# Modelling marine larval dispersal: a cautionary deep-sea tale for ecology and conservation

**DOI:** 10.1101/599696

**Authors:** Rebecca E. Ross, W. Alex M. Nimmo-Smith, Ricardo Torres, Kerry L. Howell

## Abstract

Larval dispersal data are increasingly sought after in ecology and marine conservation, the latter often requiring information under time limited circumstances. Basic estimates of dispersal are often used in these situations acknowledging their oversimplified nature. Larval dispersal models (LDMs) are now becoming more popular and may be a tempting way of refining predictions, but prior to targeted groundtruthing their predictions are of unknown worth. This case study uses deep-sea LDMs to compare predictions of dispersal. Two LDMs driven by different example hydrodynamic models are compared, along with an informed estimate based on mean current speed and planktonic larval duration (PLD) to provide insight into predictive variability. LDMs were found to be more conservative in dispersal distance than an estimate. This difference increased with PLD which may result in a bigger disparity for deep-sea species predictions. Although LDMs were more spatially targeted than an estimate, the two LDM predictions were also significantly different from each other and would result in contrasting advice for marine conservation. These results show a greater potential for model variability than previously appreciated by ecologists and strongly advocates groundtruthing predictions before use in management. Advice is offered for improved model selection and interpretation of predictions.

## Introduction

Larval dispersal is an important ecological process. Many benthic animals rely upon this phase as their only means to colonise a new area, making the process pivotal in individual survival as well as in population dynamics and persistence.

Existing global efforts to establish networks of Marine Protected Areas (MPAs) are hampered without knowledge of larval dispersal. An effective self-sustaining network needs each MPA to supply larvae to both itself and another for protected populations to persist [1] – something that will only be achieved by chance without dispersal data to base informed decisions upon. It is therefore imperative that we gather information on larval dispersal as soon as possible.

The most basic way to fulfil this need is to estimate larval dispersal using a distance /speed /time calculation based on average current speeds and planktonic larval durations (PLDs). This technique, hereafter termed “an estimate”, while highly simplistic, takes very little time, money, effort, or expertise to produce. Consequently, estimates have been used both in ecology (e.g. [2]) and conservation (e.g. [3]), although always acknowledging their oversimplified nature and the need for more detailed study.

Among the more advanced methods that exist for identifying dispersal patterns [4], larval dispersal models (LDMs) are gaining popularity in ecology and conservation (e.g. [5,6]). An LDM is a simulation of dispersal driven by a numerical hydrodynamic model to produce maps predicting which populations may be linked. As a simulation it doesn’t require expensive and difficult to obtain biological samples beyond knowing initial positional information, but it can integrate other biological data (e.g. larval behaviour, mortality, or buoyancy) should it be available (see [7]). Furthermore, the LDM can later be assessed and improved by groundtruthing with other sample-requiring methods should they become available (e.g. geochemical tracers and population genetics [4]). The ability to “provide an answer now” without requiring the time, money, and effort for additional sampling makes the LDM method particularly attractive for marine conservation’s urgent needs, especially in the deep sea [8].

However, it is well acknowledged that despite the specialist skills needed to produce an LDM, their quality and accuracy may be highly variable. Poor bathymetry, temporal and spatial averaging, a lack of sub-mesoscale processes, and unknown or estimated biological parameters can all add to the error included within an LDM [9]. The true extent of such (often unavoidable) predictive inaccuracies will always remain elusive until groundtruthing and validation can take place: essential steps in any modelling process.

Once groundtruthed, the worth of these models can be quantified, but there remains a question as to how useful un-groundtruthed LDM predictions are, and whether they should be used in preliminary management decisions? If the errors in such un-groundtruthed predictions may be large, then perhaps the crude, but fast and less expertise demanding, back-of-the-envelope estimates may be just as useful.

Shanks [10] did examine the difference between estimated and modelled predictions of dispersal distance while exploring the influence of PLD on dispersal. He found the estimate to be the least conservative prediction (an overestimate), with an LDM being up to an order of magnitude more conservative. However, the LDM also overestimated the predicted distance of dispersal when compared with those approximated from genetic data.

Shanks’s study focused on shallow-water and coastal species which are concentrated in areas of arguably more complex hydrodynamics and faster current speeds than the deep-sea. There is therefore potential for a greater similarity between estimated and modelled dispersal predictions if a similar study were focussed in deep-water.

When assessing the stability of model predictions without new sampled validation data, one approach often used in other ecological modelling disciplines, is a model comparison (e.g. [11, 12]).

It stands to reason that if all different models are trying to represent reality, there should be some similarity in their predictions, provided that their assumptions are suited to the task at hand. Exploring the differences and similarities between models promotes a greater understanding of which variables control predictions and where previously unexplored sources of error may lie.

As an ecologist running an LDM, the selection of a hydrodynamic model to power simulations is the most difficult choice to make. The huge number of models available is testimony to the variation in how they are set up – with different spatial and temporal averaging, target areas, target processes, and numerical solutions. Furthermore, each model is often supplied as source code and customised by the user so any one model name (e.g. POLCOMS, NEMO, MITgcm, ROMS) may represent a family of models where each individual iteration has been tailored to a different purpose. So it is easy to see why hydrodynamic models can appear as a black box to ecologists looking to utilise one as part of an LDM.

Despite the glut of options, model choice will be restricted first by study location and finding suitable parameterisation (e.g. see advice from [9, 13, 14]), but also by access (e.g. proprietary issues). Deep-sea studies, for example, due to the distance from shore and large spatial scales, are likely to be limited to global circulation models (GCMs), shelf models, and occasional custom build models from local observations (which carry their own limitations, see [14]).

At the end of the model selection process you may be faced with only a couple of imperfect but differently (potentially) suitable models that are hard to choose between. Allowing for parameterisation differences, a comparison of the dispersal predictions obtained from two such hydrodynamic models must logically display some difference. The question is whether the difference is negligible, and therefore potentially cross-validating, or substantial, making groundtruthing absolutely necessary before either model prediction has value.

The need to source additional data to confirm or reject model predictive ability should be considerec mandatory regardless of the result, but if models are found to broadly agree they would provide a first level of validation for each other and therefore allow meaningful research output before additional (in the deep sea, potentially considerable) groundtruthing costs are outlaid.

This study will therefore investigate:

1. The difference between estimated and modelled predictions of larval dispersal in the deep sea, and
2. The difference between the predictions of LDMs driven by two different hydrodynamic models, each selected as potentially suited to larval dispersal simulations in the study area.

By doing so this study aims to approximate the value of an un-validated LDM in relation to an estimate and discover whether a multi-modelling approach has the potential for cross-validating reassurance in predictions prior to formal groundtruthing. Note that this study does not offer a formal validation or criticism of either model, nor does it seek to recommend one over the other, instead it aims to highlight the difference and similarity between two example models to offer insight relevant to understanding the importance of model choice and the value of modelled outputs.

The results of this study should be beneficial to both ecologists and marine managers in all marine settings. The hope is to better inform those looking to use LDMs in the future and to enable a responsible interpretation of their predictions.

## Methods

### Study Area

This study was conducted in the NE Atlantic in offshore deep water west of the UK and Ireland (Fig. 1). The Rockall Trough is one of the best studied areas of deep-sea in the world, providing historic datasets for at least a qualitative groundtruthing of predictions [15, 16]. Arguably this area is not typical of deep sea due to the rapid changing bathymetry in the presence of banks and seamounts; something which can cause greater uncertainty in hydrodynamic model predictions than a flat abyssal plain [9]. This could however make for a fairer comparison to complex shallow water and coastal hydrodynamics and also promotes a greater similarity to estimate predictions which represent a null model of maximal uncertainty and spreading of larvae.

**FIGURE 1.**
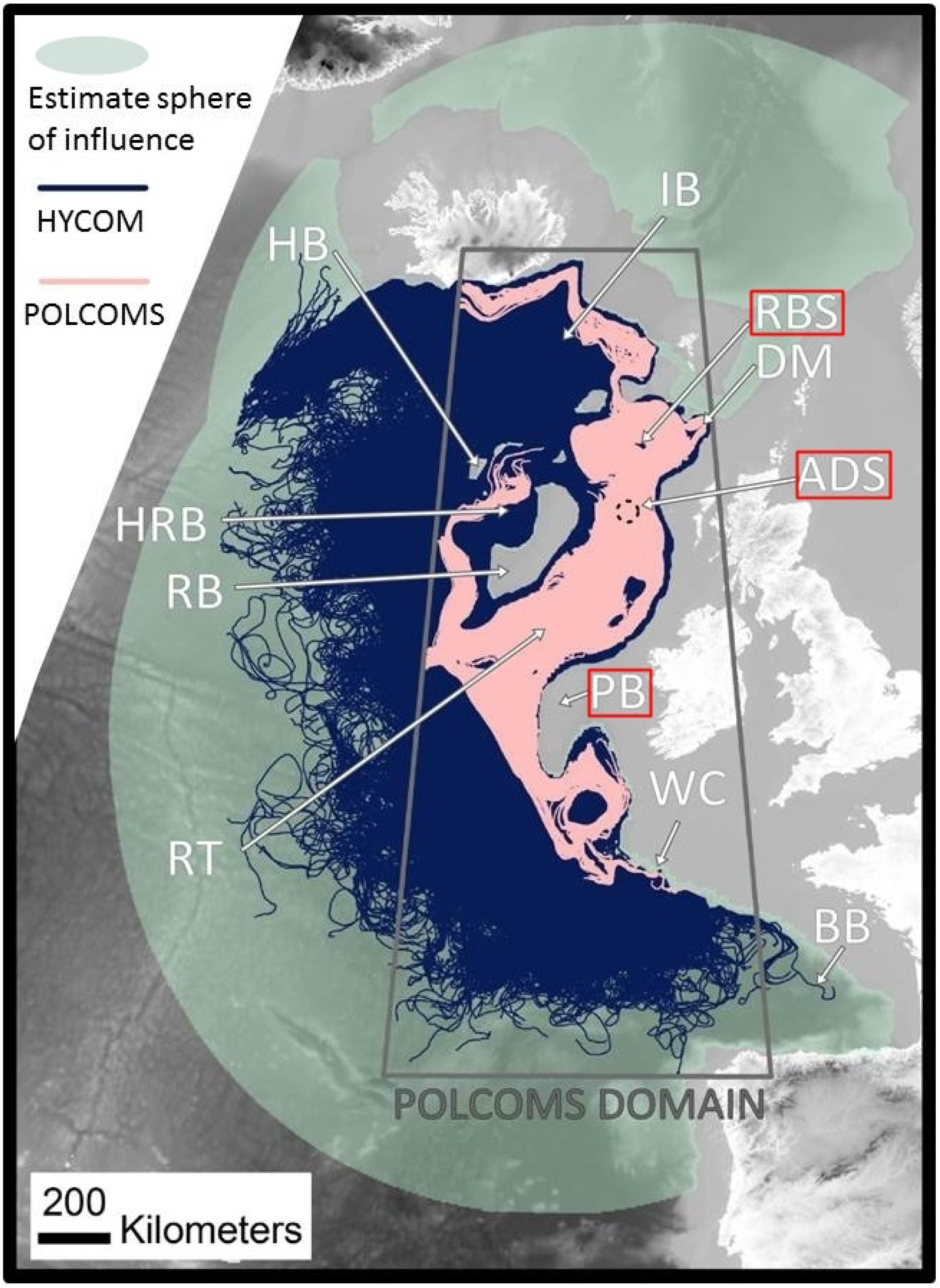
Plot of study area and all simulated tracks in the HYCOM and POLCOMS larval dispersal models relative to the sphere of influence predictions of an average current speed based estimate (0.1 m s^−1^). The grey box delineates the domain of the POLCOMS model: tracks simulated by the POLCOMS model are unable to exit this area. Larvae were released from Rosemary Bank (RBS), Anton Dohrn Seamount (ADS) and Porcupine Bank (PB) (highlighted with red boxes). Features of topography mentioned in the text are labelled as follows: Iceland Basin (IB), Hatton Bank (HB), Darwin Mounds (DM), Hatton Rockall Basin (HRB), Rockall Bank (RB), Whittard Canyon (WC), Bay of Biscay (BB). (This map was created in ArcGIS 10.3 (http://www.esri.com) with GEBCO 30 arc-second topography, available from www.gebco.net, and projected Albers Equal Area Conic with modified standard parallels and meridian (sp 1 = 46 °N, sp 2 = 61 °N, m = 13 °W)).

### Estimate calculation

This deep-sea case study relates findings to a figure published in McClain & Hardy [2]. The figure, notably with a caption full of caveats, displays potential larval dispersal distances of deep-sea fauna based on two different potential deep-sea averaged current speeds derived from Havenhand *et al.* [17]. This study will use the lower averaged current speed (0.1 m s^−1^) as the estimate, after Ellett, Edwards & Bowers [15] who cite a vector-averaged current speed (over 15 day periods between 1975-1982) of 0.1-0.2 m s^−1^ in the study region, as recorded in the vicinity of Anton Dohrn Seamount.

### LDMs

Two LDMs were run in this study, each consisting of a single particle simulator paired with one of the two hydrodynamic models; additional details on all model algorithms and parameterisations are available in Supplementary Material S1.

### Particle simulator: Connectivity Modeling System (CMS)

The CMS was used as the particle simulator (hereafter “simulator”) for both LDMs. There are many types of simulator available, but, without in-depth numerical modelling expertise, ecologists are likely to be limited to the use of offline simulators paired with the outputs from a hydrodynamic model (see Hilãrio *et al.’s* [8] supplementary table for a list of offline simulators and their compatibilities). The CMS is one such offline simulator. It is both freely available and designed especially with LDM in mind (https://github.com/beatrixparis/connectivity-modeling-system, v 1.1, [18]). This simulator has shown success in recent estimates of species connectivity [19–21] as well as driving investigations of abyssal hydrodynamic transport [22] among other studies.

While it is easy to integrate biological data, this study uses the simulator it in its simplest configuration simulating passive dispersal for the cleanest comparison. An hourly particle tracing timestep was used as decided by model sensitivity testing [23], and positional outputs were recorded daily.

### Hydrodynamic Model 1: POLCOMS

POLCOMS is a shelf and coastal model used in UK and Irish waters. This version comes from Plymouth Marine Laboratory, UK. It was previously used by the UK Met Office in weather forecasting – a fact which might recommend it above other models in this area [24, 25] and has been extensively validated over the UK surrounding waters [26]. The 1/6 ° x 1/9 ° (c. 12 km^2^) resolution offers an eddy-resolving solution, however it can only capture major eddies (c. 64 km in size based on needing six or more data points to adequately resolve an eddy [27]) making this the coarser of the two models trialled. The model was run with 40 terrain following depth layers (sigma-levels) although outputs were interpolated to a z-level format (a list of set depth levels) using Matlab (v.R2013a) in order to make them compatible with the CMS. POLCOMS has been used in several dispersal studies to date (e.g. [28, 29]).

### Hydrodynamic model 2: HYCOM

HYCOM is a freely available global hydrodynamic model developed by the US Navy (www.hycom.org, [30]). It is uniquely set up to use a hybrid of water mass following, terrain following, and depth specific vertical layers, changing with the underlying topography, which may make it well suited to deep-sea studies. The outputs, however, are in the z-level format required by the CMS (but may lose some of the hybrid grid details in the reformatting). The 1/12 ° resolution (c. 8 km x 4 km in the study area), allows smaller eddies (c. 48 km wide) to be captured than in POLCOMS, although this is still coarse relative to reality. The global nature of HYCOM may be an upside for wide-ranging studies but is also a downside as the validation of the model was performed on a global scale and it may therefore not validate so well on a local scale [14]. HYCOM has already been used in multiple dispersal studies [30–33], including in the deep-sea [34, 20, 23, 35].

### Larval releases

“Larvae” were released from three locations in the Rockall Trough in order to access different current regimes in the area: Rosemary Bank in the north, Anton Dohrn Seamount in the centre, and Porcupine Bank in the south (see Supplementary Material S2 for exact positions). Releases were made from 16 positions per depth band from four depths (700 m; 1,000 m; 1,300 m; 1,500 m).

Releases were made daily for 366 days from 4^th^ January 2003 – 4^th^ January 2004. All particles were tracked for 270 days in line with McClain & Hardy [2], although daily positional outputs allow subsetting of this PLD.

Simulations were run without additional diffusivity parameters as this would complicate interpretation of the difference in hydrodynamic model instruction, would be a different setting for each model, and would be subjectively chosen as a nest-wide parameter. This decision is in line with the study undertaken by Shanks [36] in comparison to Siegel *et al.* [37]. As a result of excluding diffusivity only one particle is released per day as simultaneous releases will follow identical tracks. N.B., We do advocate adding diffusivity parameters in larval dispersal studies that are focussed on ecological questions (rather than the aim of this study – to compare the difference in instructions given by different hydrodynamic models). Indeed, diffusion can be of greater importance than advection [38], so this parameter should be considered carefully.

Neither of this study’s hydrodynamic models supply vertical velocity fields (w) due to their large spatial domains so simulations in this case are effectively 2-dimensional.

### Analysis

In order to perform a comparison meaningful to ecologists and marine managers, both distance and spatial predictions were analysed. Ecologists often examine dispersal kernels and the potential distance of larval dispersal (terrestrial examples [39–41]; and marine examples [2, 37, 4, 42]), while marine managers may require more spatially explicit descriptions examining whether Location X is connected to Location Y [43–45].

### Distance comparisons

Distance comparisons were illustrated by converting larval fates into effective dispersal kernels. CMS outputs consisting of daily positions of each simulated particle were converted into straight line distance (SLD) from source, per day, in Matlab (version R2013a) using the Haversine formula to account for earth curvature. A median SLD per day was then calculated for each model, as well as per depth per model, and associated quartiles. The result was plotted against the average speed 0.1 m s^−1^ line in the same format as the McClain & Hardy [2] figure for ease of comparison.

The difference between the median SLD per day per model was tested using a negative binomial GLM accounting for depth and location. An analysis was undertaken for the full 270 day time frame, with noted reference points at 35 days and 69 days tracking which were discerned by Hilãrio *et al.* [8] as the median and 75 % quartile PLDs of all deep-sea and eurybathic species where PLD is known (n=92 species).

### Spatial comparisons

Larval fates were mapped to offer a means of qualitative and quantitative spatial comparison. The maps were created in ArcGIS (version 10.1) using an Albers Equal Area Conic Projection with modified standard parallels (46 °N, 61 °N). A grid of constant 4 km^2^ cell size (approximately half the HYCOM model resolution) was applied across the POLCOMS domain (see Fig 1; the prediction area with the most restrictive boundaries). For each depth band, grid cells occupied by topography were removed resulting in the 2D maximal possible area of occupancy.

The estimate was mapped as a sphere of influence buffer zone prediction with radius equal to the average predicted dispersal distance. The major limitation of an estimate prediction is that it cannot easily be extrapolated into a probabilistic spatial prediction without a method to quantify the error caused by assuming a constant current direction. All areas within the buffer zone were therefore considered as a presence-only record. The estimate prediction was therefore always equivalent to the 2D maximal possible area of occupancy.

The prediction from each LDM was mapped as a percentage track density per grid cell occupancy in order to provide a spatial “heat map” of dispersal. Track density values were used for the quantitative LDM comparison, while the estimate versus model comparison required a binary (presence only) comparison of occupied cells.

A cumulative cell by cell linear correlation coefficient computed in R offers a single correlation value as representative of the comparison between each prediction (a raster correlation). This was performed per release location, per depth, and summarised as an average correlation between models.

Additional qualitative assessments offer real world interpretations of potential connectivity between sites. These are relevant to how usefully similar predictions may be to marine management and conservation.

## Results

### LDMs vs Estimate

Both in terms of distance and spatial dispersal patterns, the estimate prediction was the least conservative and specific, with both modelled predictions being considerably more retentive and spatially targeted (Fig. 1 and Fig. 2a,b).

**FIGURE 2.**
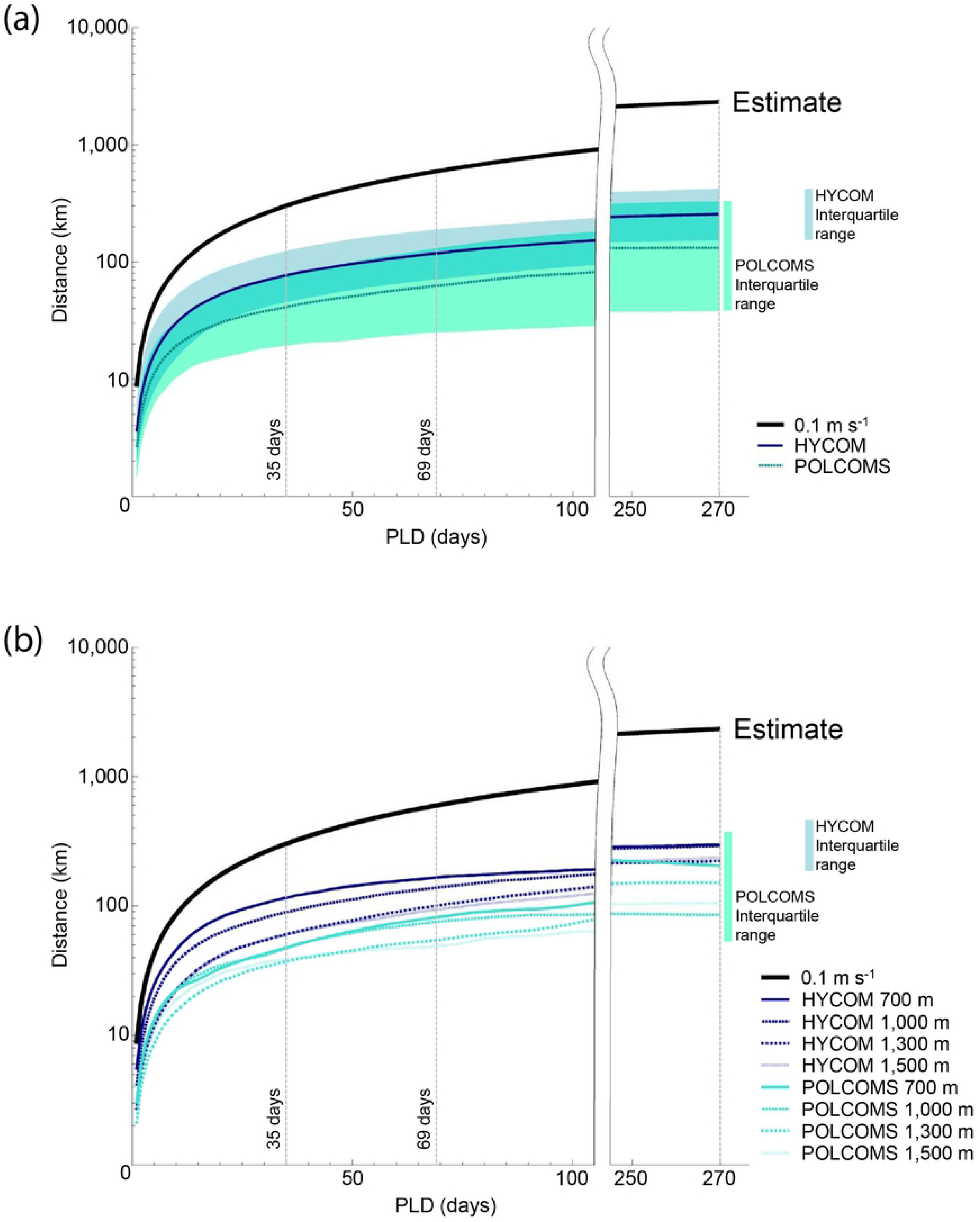
Results plots of median dispersal distance over time for the estimate (after McClain & Hardy^40^), and model predictions. (A) Median values per model with shaded interquartile ranges inclusive of overlap. (B) Median values per depth band per model. Reference PLDs are highlighted in line with Hilãrio et al. [8] and the PLDs representative of 50 % (35 days) and 75 % (69 days) of all known deep-sea animals.

### Distance

Plots of median dispersal distance over time show a clear difference between the two LDM predictions and estimate (Fig. 2a) both when considered together (ANOVA p < 0.0001, F (2,809) = 641.5) and individually (post-hoc Tukey HSD p < 0.0001). The estimate offered the least conservative dispersal distances, being almost double the LDM predicted dispersal distance from day one, scaling to an average five-fold increase at 35 days, almost seven-fold at 69 days, and twelve-fold at the full 270 days tracking.

### Spatial – correlation

The HYCOM LDM spatial predictions were the most similar to the estimate, although the similarity was still less than 0.5 (0.44 across all depths and locations) (Table 1). The correlation between the estimate and POLCOMS LDM was very weak averaging 0.17 across all depths and locations. Across all depths and locations the correlation between estimated and modelled spatial extents was max. 0. 67 (HYCOM LDM, Porcupine Bank, 700 m simulations), and min. 0.06 (POLCOMS LDM, Rosemary Bank, 1,500 m simulations).

**TABLE 1.** Linear correlation coefficients between predictions provide quantitative spatial comparisons between LDM and estimate predictions. Binary correlations are between presence only grids, while the track density correlation between LDMs are sensitive to the full spatial spread as well as the locations of “dispersal highways”. Minimum values are highlighted in *<u>underlined italics</u>*, and maximum values in **bold**.

|  | Depth | ROSEMARY<br>BANK | ANTON<br>DOHRN | PORCUPINE<br>BANK | All depths<br>& sites |
| --- | --- | --- | --- | --- | --- |
| POLCOMS<br>LDM Vs<br>Estimate<br>(binary) | 700 m | 0.22 | 0.11 | 0.24 | 0.17 |
|  | 1,000 m | 0.19 | 0.13 | **0.25** |  |
|  | 1,300 m | 0.16 | 0.15 | 0.21 |  |
|  | 1,500 m | <u>0.06</u> | 0.1 | 0.24 |  |
| HYCOM<br>LDM Vs<br>Estimate<br>(binary) | 700 m | 0.4 | 0.44 | **0.67** | 0.44 |
|  | 1,000 m | 0.39 | 0.41 | 0.6 |  |
|  | 1,300 m | 0.28 | 0.29 | 0.59 |  |
|  | 1,500 m | <u>0.24</u> | 0.25 | 0.66 |  |
| POLCOMS<br>Vs HYCOM<br>LDMs<br>(binary) | 700 m | **0.52** | 0.25 | 0.34 | 0.36 |
|  | 1,000 m | 0.42 | 0.31 | 0.41 |  |
|  | 1,300 m | 0.49 | 0.46 | 0.35 |  |
|  | 1,500 m | <u>0.18</u> | 0.31 | 0.32 |  |
| POLCOMS<br>Vs HYCOM<br>LDMs<br>(density) | 700 m | 0.23 | 0.39 | **0.46** | 0.35 |
|  | 1,000 m | **0.46** | **0.46** | 0.37 |  |
|  | 1,300 m | <u>0.09</u> | 0.35 | 0.45 |  |
|  | 1,500 m | 0.19 | 0.33 | 0.35 |  |

### Spatial – qualitative

Qualitative spatial comparisons between estimate and LDM predictions further emphasise their difference. For example, neither LDM suggests connections as far as the Spanish/Portuguese continental shelf while the estimate would (Fig. 1).

The difference between the estimate and the POLCOMS LDM is the most pronounced. Even though the POLCOMS domain is the most restricted, there is a large area within that domain that remains untouched by POLCOMS dispersal pathways. For example, none of the POLCOMS releases connect to much of the Hatton Rockall Basin, the north side of Hatton Bank, or south beyond the Whittard Canyon in the Bay of Biscay (Fig. 1).

Maps per model, location, and depth (Fig. 3–5) allow visualisation of more detailed comparisons. For example, Porcupine Bank simulations (Fig. 5) suggesting no connection to Rosemary Bank at 1,300 m and 1,500 m in either model, while the estimate would comfortably make that distance. This would make a difference to a marine manager who might want to know whether known fauna at 1,500 m depth can reach a protected area at Rosemary Bank: an estimate would say ‘yes’, and an LDM would say ‘no’.

**FIGURES 3-5.**
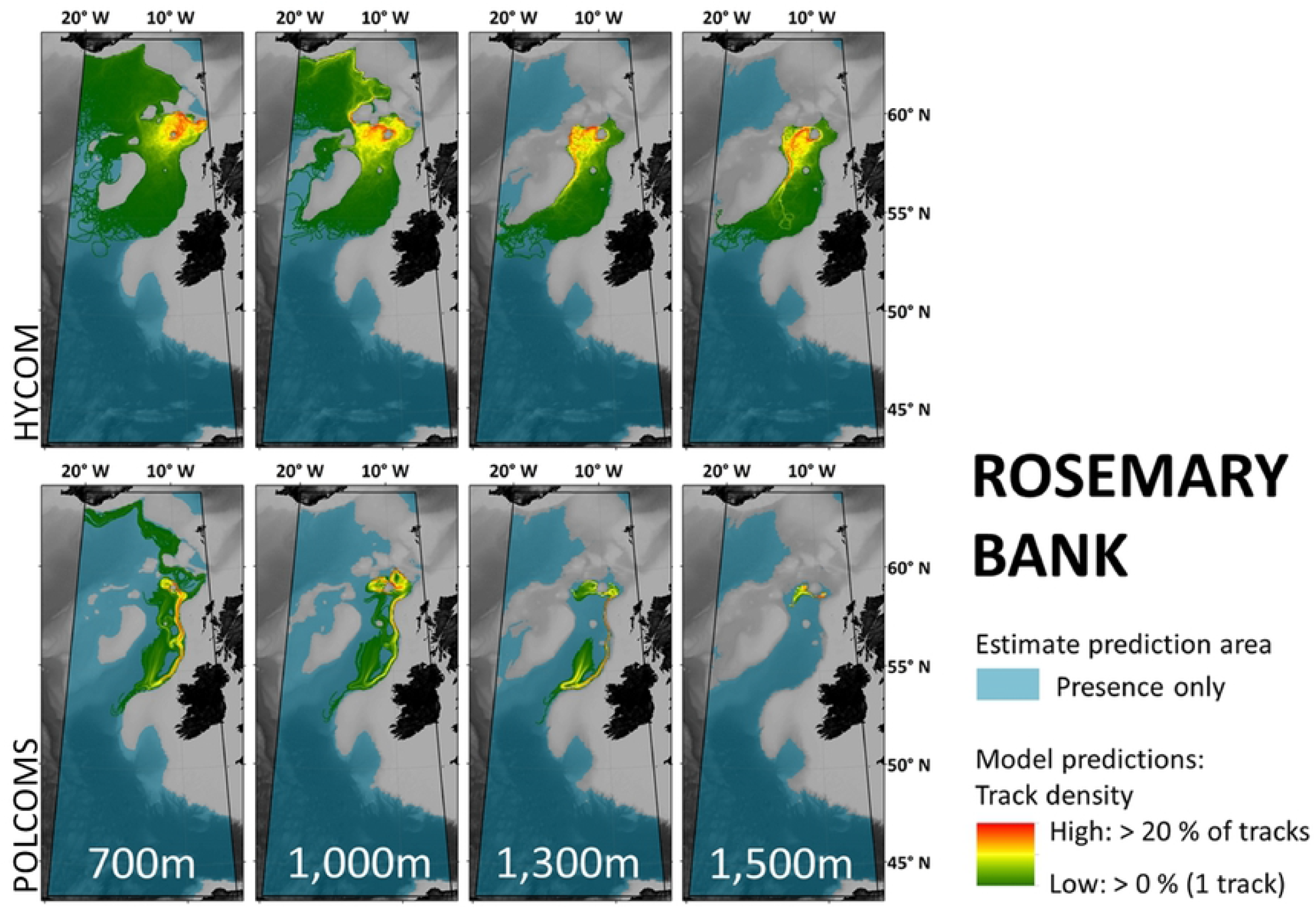
Maps per depth band of predicted larval dispersal as simulated from Rosemary Bank (Fig. 3), Anton Dohrn Seamount (Fig. 4) and Porcupine Bank (Fig. 5). Simulations from HYCOM and POLCOMS models are displayed as track densities delineating between occasional and persistent pathways of dispersal. The estimate prediction area fills the POLCOMS domain but due to the 2D nature of simulations excludes areas of raised topography. Spatial correlations were conducted comparing the extent of modelled and estimated predictions and the extent and density information of each modelled prediction (Table 1). (All maps were created in ArcGIS 10.3 (http://www.esri.com) with GEBCO 30 arcsecond topography, available from www.gebco.net, and projected Albers Equal Area Conic with modified standard parallels and meridian (sp 1 = 46 °N, sp 2 = 61 °N, m = 13 °W)).

**FIGURES 3-5.**
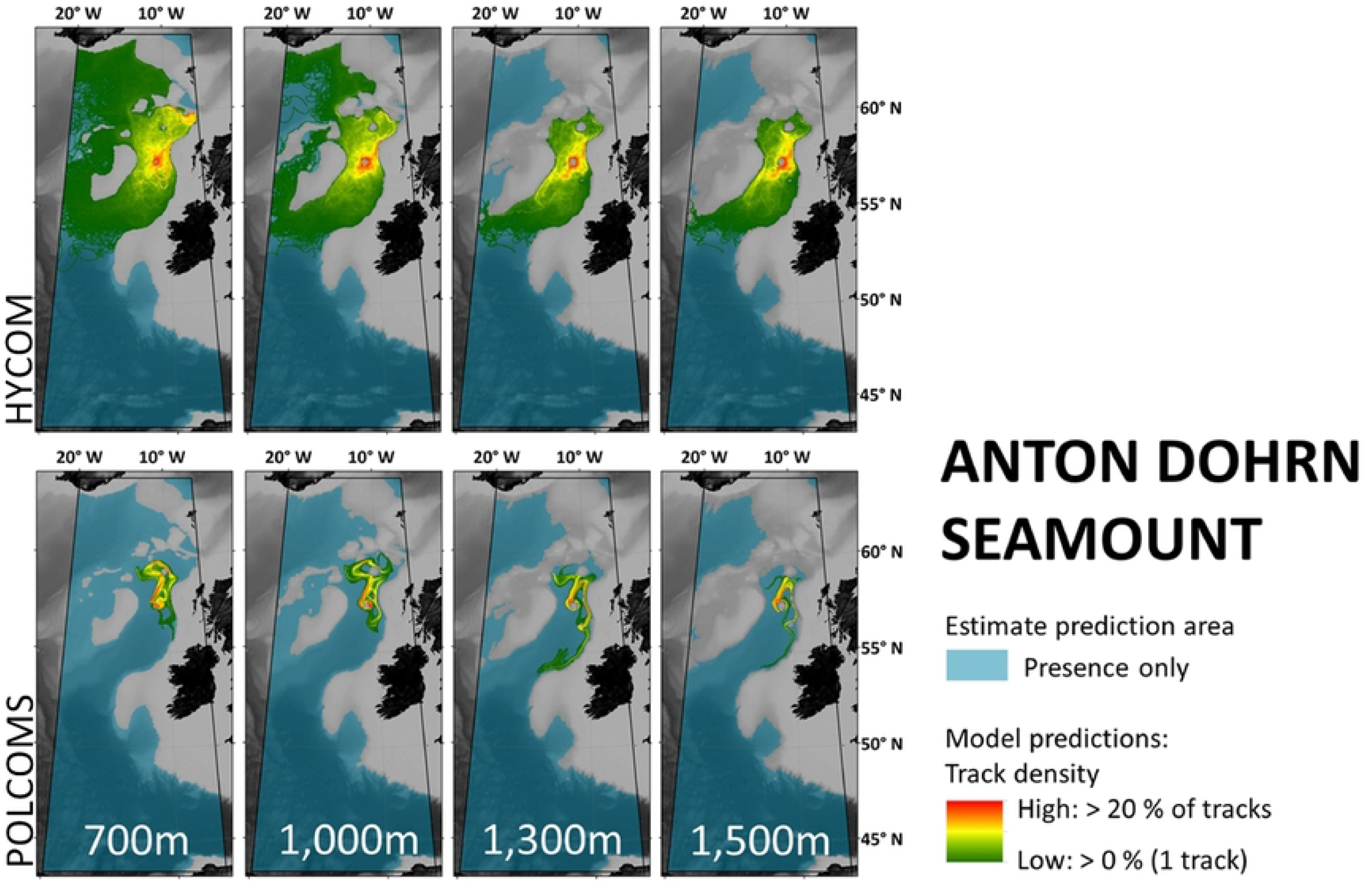
Maps per depth band of predicted larval dispersal as simulated from Rosemary Bank (Fig. 3), Anton Dohrn Seamount (Fig. 4) and Porcupine Bank (Fig. 5). Simulations from HYCOM and POLCOMS models are displayed as track densities delineating between occasional and persistent pathways of dispersal. The estimate prediction area fills the POLCOMS domain but due to the 2D nature of simulations excludes areas of raised topography. Spatial correlations were conducted comparing the extent of modelled and estimated predictions and the extent and density information of each modelled prediction (Table 1). (All maps were created in ArcGIS 10.3 (http://www.esri.com) with GEBCO 30 arcsecond topography, available from www.gebco.net, and projected Albers Equal Area Conic with modified standard parallels and meridian (sp 1 = 46 °N, sp 2 = 61 °N, m = 13 °W)).

**FIGURES 3-5.**
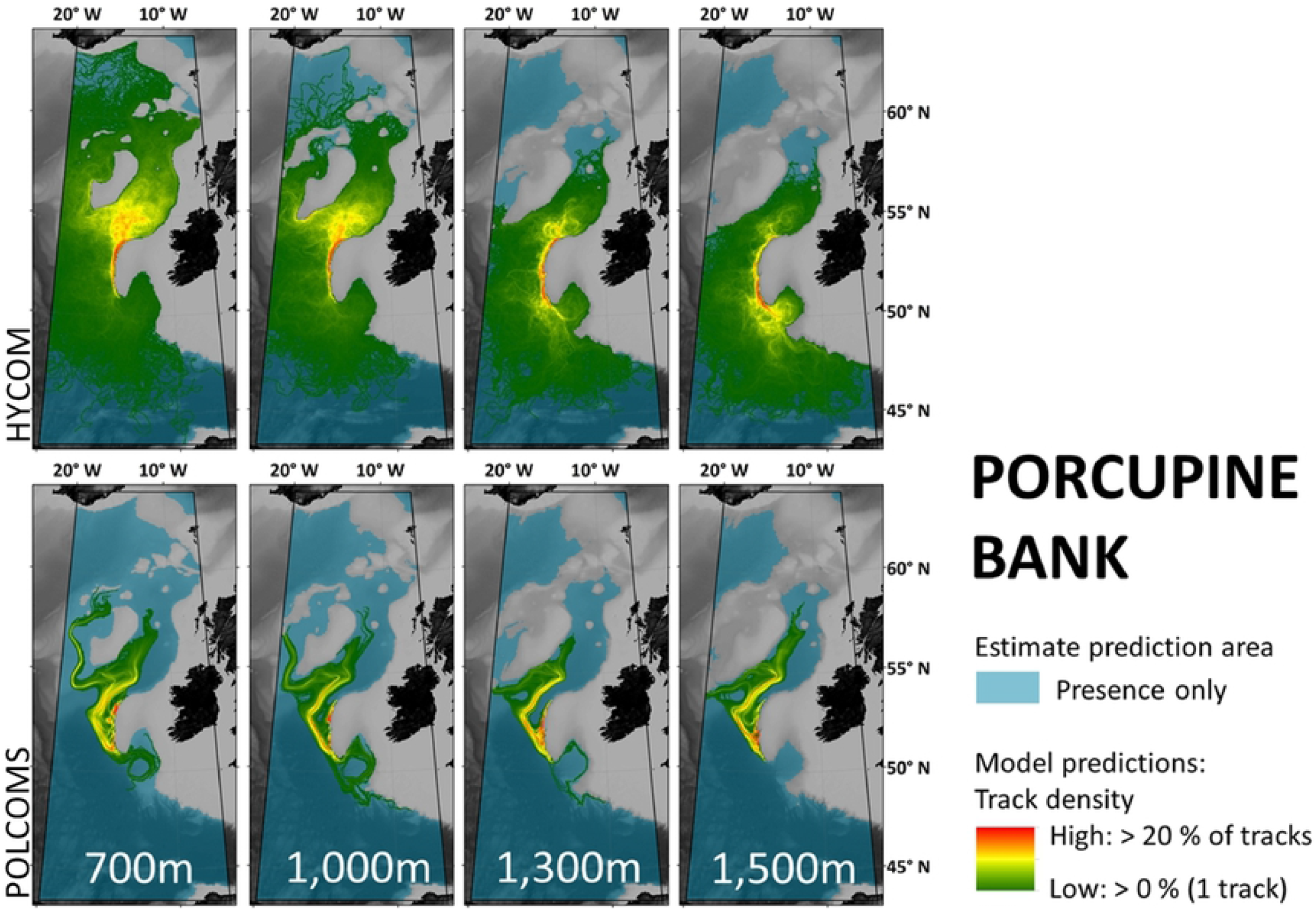
Maps per depth band of predicted larval dispersal as simulated from Rosemary Bank (Fig. 3), Anton Dohrn Seamount (Fig. 4) and Porcupine Bank (Fig. 5). Simulations from HYCOM and POLCOMS models are displayed as track densities delineating between occasional and persistent pathways of dispersal. The estimate prediction area fills the POLCOMS domain but due to the 2D nature of simulations excludes areas of raised topography. Spatial correlations were conducted comparing the extent of modelled and estimated predictions and the extent and density information of each modelled prediction (Table 1). (All maps were created in ArcGIS 10.3 (http://www.esri.com) with GEBCO 30 arcsecond topography, available from www.gebco.net, and projected Albers Equal Area Conic with modified standard parallels and meridian (sp 1 = 46 °N, sp 2 = 61 °N, m = 13 °W)).

The low correlation between the estimate and POLCOMS LDM in particular can be exemplified by Fig. 3. Here the estimate might expect connections between Rosemary Bank and anywhere in the Rockall Trough and Bay of Biscay to the south, while the POLCOMS LDM suggests larvae may not even reach neighbouring Rockall Bank in the west at any depth. Therefore, a marine manager asking whether there would be a dispersal connection between Rosemary Bank and Anton Dohrn Seamount would be told ‘yes’ from an estimate, and ‘no’ from a POLCOMS LDM.

## HYCOM LDM vs. POLCOMS LDM

Generally, the two LDMs tested in this study give notably different predictions of dispersal, displaying differences in distance, spread, and, in some cases, direction of travel.

### Distance

Figure 2a shows the lower median dispersal distances and much larger interquartile range of the POLCOMS predictions when compared to HYCOM. An ANOVA comparing the two LDM’s median distances per day confirms this difference (p < 0.0001, F (1,538) = 276.8). Plots of median dispersal distance per depth (Fig. 2b) demonstrate that the shallowest (most dispersive) simulations in the POLCOMS model on average travel less far than the deepest (least dispersive) simulations in the HYCOM model.

### Spatial – correlation

The correlation between the track density maps of each LDM was generally low (Table 1, bottom section). The maximum correlation between POLCOMS and HYCOM simulations was 0.46 (Rosemary Bank 1,000 m; Anton Dohrn 1,000 m; and Porcupine Bank 700 m simulations), minimum 0.09 (Rosemary Bank 1,300 m simulations) and the average across all depths and locations only 0.35. NB This can be compared to the presence-only correlations which are also shown in Table 1 and are still low: max 0.52 (Rosemary Bank 700 m), min 0.18 (Rosemary Bank 1,500 m), av. 0.36.

### Spatial – qualitative

Generally, Rosemary Bank simulations were the most dissimilar (Fig. 3). For example, while the 1,500 m Rosemary Bank simulations in HYCOM suggest connection southwards to most of the eastern flank of Rockall Bank, POLCOMS predicts a relatively small dispersal range suggesting there may be no connection to Rockall Bank at all.

Of most concern is when the two models disagree in the direction of dispersal. In Rosemary Bank 1, 300 m simulations, HYCOM show the “highways” of high track density extending west down the eastern flank of Rockall Bank, while POLCOMS extends down the east of the Rockall Trough following the continental slope. Indeed in 1,000 m Rosemary Bank simulations, HYCOM larvae travel North, while POLCOMS larvae travel South.

By contrast, the results from Anton Dohrn Seamount (Fig. 4) are more similar, with all “highways” generally extending north-east towards Rosemary Bank in both HYCOM and POLCOMS simulations. Yet if a marine manager were to ask whether larvae from Anton Dohrn reach the Darwin Mounds to the north-east, HYCOM would say ‘yes’ and POLCOMS would say ‘no’.

Simulations from Porcupine Bank (Fig. 5) might indicate a broad agreement that larvae will eventually reach the southern Rockall Bank, but the less direct “highways” in the POLCOMS model might reduce chances of larvae getting that far.

## Discussion

This study explored the value of larval dispersal predictions from LDMs by considering two questions.

### 1) Will LDMs give a notably different result to an (informed) estimate?

Our results agree with Shanks [10] and suggest that yes, there can be a large difference in the predicted distance, area, and specificity of estimated and modelled dispersal patterns. There could therefore be a distinct advantage in going to the effort of modelling predictions, provided that models are shown to adequately approximate realistic distances better than the estimate.

This study may also suggest that for deep-sea species the difference may be even more pronounced than in shallow water. As the difference in predicted distance of dispersal increased exponentially with tracking time (up to a 12-fold difference), species with longer PLDs, such as those from the deep sea [8], may show even greater disparity between estimated and modelled predictions.

However, there may have been better congruity between estimated and modelled predictions if simulations were undertaken in the open ocean. If the estimate were similar to the modelled predictions this would suggest that the model simulates currents with fairly straight trajectories and constant speeds: something more likely to occur on a relatively featureless abyssal plain at 5,000 m. The complex topography of the Rockall Trough induces a lot of mesoscale activity [46] which likely promotes greater local retention, and therefore difference in modelled predictions. Alternatively estimates could be made to better approximate what the models include; either by following the topography to account for the distance added by including depth, as is the case in 3D models, or by at least accounting for topographic barriers at the simulated depth, which would be a closer approximation to this study’s 2D simulations.

### 2) Are two LDMs, driven by different purpose-selected hydrodynamic models, cross-validating and therefore of some value prior to targeted groundtruthing?

Broadly, while some local comparisons may be cross-validating (e.g. in Anton Dohrn simulations), in this study the different hydrodynamic models gave very different predictions. Indeed, the variability in the predictions suggests that the potential for error within LDMs may be larger than previously recognised in ecology and conservation – a result that cannot be apparent when modelling with only one hydrodynamic model. This result emphasises that in all areas where the models disagree, there can be no trusted consensus until targeted groundtruthing takes place, and that the un-groundtruthed LDM outputs must not act as a basis for decision-makers before a groundtruthed consensus can be reached.

### Model differences

While we are not going to provide any criticism or endorsement for either model, we can offer some limited analysis and advice to aid model selection and interpretation in the future. In this case, there are three hydrodynamic model parameters that are worth highlighting which may account for the differences in LDM predictions.

First, there are differences in the **scales of validation** between the models, but also in the relevance of these validations for dispersal modelling purposes. Despite both models being published and validated, HYCOM was assessed on a global scale and therefore may potentially be less locally reliable. However, neither model was validated for the purpose of larval dispersal modelling which may place greater weight on, for example, current directions and strengths than heat exchange and mixed-layer behaviour. This makes it hard to use a model’s published validation status to judge whether the model is fit for purpose and recommends that study specific validation is vital, starting with a comparison to observational oceanography in the area [33]. In this study, for example, the southward trajectories of POLCOMS larvae down the eastern side of the Rockall Trough from Rosemary bank at 700 m – 1,300 m (Fig. 3) are contrary to the observations of northward transport down to 1,000 m, and below that southward transport down the western side of the Trough [16, 47]. However current speeds simulated in each model are different, with velocities in HYCOM being twice those in POLCOMS, although both fall within the range of observed current speeds recorded in the shelf edge current (10-21 cm s^−1^) [48]. Note that both models will suffer from many other errors including (but not limited to) currents that are too fast due to the exclusion of tides [49], coarse bathymetry that may exclude hydrographically influential features [50], and no representation of possible benthic storms which may divert dispersal pathways [51]. Only targeted groundtruthing can quantify the error margins and clarify whether one model is more representative than the other for this purpose, and indeed they may each prove to have areas of accuracy at different depths or locations [33].

Second, **Spatial and temporal resolution** has been shown to make a great difference in whether a model represents realistic trajectories or not. Putman & He [52] advocate using the highest resolution model you can find, summarising that model choice must aim to preserve physical processes on the scale tens of kilometres and days (respectively). Although both models may comply with this broad advice, HYCOM is still more highly resolved than POLCOMS. In this study area, while the major eddies may be over 100 km in diameter [53], there are still some influential semipermanent features of 50-60 km in diameter [54, 55]. Given a rule-of-thumb that six data points are required to make an eddy [27], POLCOMS may omit these smaller eddies (min. eddy size of ~64 km in diameter), while HYCOM may be capable of capturing them (min. ~48 km in diameter). This difference in horizontal resolution may account for some of the difference in trajectory direction and tortuosity between the models. Consequently, we recommend that minimum resolution choice could be based on the size of permanent eddies in the study region (if that information is known).

Third, and finally, algorithm **error handling** may be responsible for the more diffuse trajectories in the HYCOM model, but this is much harder to account for when choosing a model. The horizontal pressure gradient error stems from the issue of interpreting flow around steep discretized (pixelated) topography and can result in perpetuated errors throughout the water column. This issue is handled in both models but using different approaches (see Supplementary Material S1 for model approaches). A representation of this can be seen in Supplementary Material S3, where plots of current ellipses per model, per depth, can help highlight these differences: the less variable current direction and speed (smaller ellipses) and tight shelf edge current in the POLCOMS model suggests a stricter handling of these errors, resulting in a less diffusive spread of particle trajectories. Accounting for this difference during model choice is more problematic for non-numerical-oceanographers but highlighting its effect here may offer a means of recognition and inform interpretation.

### Groundtruthing

Groundtruthing should be regarded by all modellers as essential, and were the models found to be similar it would not have supplanted this necessity but could have lent some credence to modelled outputs before groundtruthing data became available. As it stands, however, the tested models could not be used interchangeably without consequence to ecological or marine management conclusions (e.g. whether Rosemary Bank was connected to south-east Rockall Bank at 1,300 m (Fig. 3)). Hence the next step must be to identify whether one model is more accurate than the other.

There have been model predictions with groundtruthing success, particularly when comparing LDMs and seascape genetics [56, 57, 21]. Once groundtruthed, an LDM could be incredibly useful across disciplines and purposes, allowing subsequent simulations in the same region to be run and trusted for multiple species provided that similar oceanographic features are important to larval fates. New species predictions would then be able to rule out hydrodynamic sources of error leaving biological components as the main areas requiring groundtruthing in the future. It is therefore advisable to build the first species LDMs in a region upon the species with the greatest amount of data available for both model parameterisation and groundtruthing. Once completed and tested, LDMs for other species can be created, safe in the knowledge that error due to hydrodynamic model choice is now quantified and controlled for.

### Advice for marine managers and ecologists

In this case, the variation in direction of dispersal makes it unwise to rely upon these predictions until they have been groundtruthed. However, were the differences only in speed and spread, these predictions may have been more useful, allowing interpretation relative to appropriate precautionary principles for the issue being considered. For example, MPA network design may wish to accommodate the most conservative predictions of dispersal to ensure that the larvae from a protected population can reach the next protected area (in this instance that would be POLCOMS predictions). Meanwhile if you were estimating invasive species spread, you may wish to default to a less conservative estimate (here HYCOM).

The variability between models also advocates the interpretation of LDM results as probabilistic (i.e. possible) rather than deterministic (i.e. true). Practically this may be translated as looking at the high density “highways of dispersal” which had some localised consensus between models, so these could be interpreted as the more likely pathways of dispersal, with lower density predictions being thought of as uncertain.

In summary, LDMs will have a place in marine ecology and conservation and offer a great improvement on informed estimates of dispersal potential, however, the hydrodynamic models they are based on can be very variable in their predictions, so should always be assumed to need some level of study-specific groundtruthing prior to relying upon predictions for management decisions and ecological theories. Utilising local oceanographic observations and model comparisons can indeed offer some basic means of quantifying the uncertainty in model predictions to improve trust, but future comparison to population genetics, geochemical isotope tracers, or study-targeted groundtruthing data must still be considered essential.

## Acknowledgements

The authors would like to thank Plymouth Marine Laboratory for supplying outputs from the POLCOMS model; Nataliya Stashchuk for assistance in interpolating POLCOMS from sigma to z-level format; Peter Mills and Antonio Rago for High Performance Computing support; Vasyl Vlasenko for his supervision; and Antony Knights, John Spicer, and Anthony Grehan for their advice.

## Supporting information

**S1 – Additional detail on model algorithms and parameterization**

Some additional descriptions of a) Connectivity Modeling System (CMS) – the particle simulator, b) POLCOMS hydrodynamic model, c) HYCOM hydrodynamic model.

**S2 – Model release locations**

Including a) HYCOM Release Locations, b) POLCOMS Release Locations, c) Why there is a difference between the release locations in each model, d) An overview map of release locations.

**S3 – Current ellipses**

Plots focussed on the shelf edge current that can demonstrate the different handling of horizontal pressure gradient errors between the two hydrodynamic models, POLCOMS and HYCOM.

